# Comprehensive profiling of antibodies against multiple infectious diseases in serum and the airway mucosa using synthetic peptide-based linear epitope microarrays

**DOI:** 10.1101/462689

**Authors:** Charles J Sande, Timothy Chege, Martin N Mutunga, Elijah Gicheru, D James Nokes, Christopher A Green, James Tuju

## Abstract

**Background:** Despite the considerable progress that has been achieved in the development of vaccines, infectious diseases continue to be the major cause of morbidity and mortality in developing countries. The first year of life remains the time of greatest risk with more than half of all deaths caused by infectious diseases in children under the age of five years occurring during this period. For most infections, antibodies are the strongest immune correlate of protective immunity and characterizing the breadth of the antibody repertoire against infectious diseases in early life is key to profiling individual disease risk.

**Methods:** In order to comprehensively profile the antibody repertoire against infectious diseases in early life, we developed a synthetic peptide-based microarray to simultaneously measure antibodies against forty one common infectious diseases. 82 empirically-validated linear B-cell epitopes were selected from the Immune Epitope Database (IEBD), expressed as synthetic peptides and printed onto microarray glass slides (microarray chip). The chip was used to simultaneously measure antigen-specific IgG and IgA in serum and mucosal samples from thirty eight infants and young children presenting to hospital with different illnesses. We also used the data from the microarray to estimate antibody decay kinetics for the five most frequently detected antigens during the first year of life.

**Results:** A synthetic peptide microarray to measure antibodies against forty one infectious diseases was successfully developed. Although the combination of antigens that were recognized were different for each child, antigens derived from Epstein-Barr virus, poliovirus, *Streptococcus pneumoniae*, plasmodium falciparum and varicella zoster were recognized by most study participants. While the combination of antigens recognized in serum was generally similar as those recognized in mucosal samples, antibodies to antigens such as herpesvirus, rubella and echinococcus were more predominant in either serum or mucosal samples. With the exception of the pneumococcus, we observed a progressive decline in serum IgG specific to all the infections above in the first six months of life. Pneumococcal IgG in serum exhibited a continuous rise in the first six months of life.

**Conclusions:** We successfully developed a synthetic peptide microarray and used it to profile the diversity of systemic and mucosal antibody against multiple infectious diseases in children. The data from the slide provide a snapshot overview of the breadth and kinetics of infectious disease antibodies early in life and opens the opportunity for conducting in-depth multi, multi-target serological studies of infectious diseases in future.

## Introduction

Susceptibility to infections in early life can be attributed to inadequate immune protection against the multiple infections to which infants and young children are constantly exposed^1^. For most infections, antibodies correlate strongly with protective immunity as demonstrated by the fact that most licenced vaccines function through the induction of functional antibodies^2^. In early life, these protective antibodies can be also derived from alternative sources including the immune response mounted against infection, passively-acquired maternal antibody^3^ and maternal breast milk4. Antibodies from different sources contribute towards an immune repertoire whose breadth defines the range of infections to which an individual is susceptible. In order to comprehensively profile the antibody repertoire against infectious diseases in early life, it is necessary to simultaneously measure antibodies against multiple antigens within a single biological sample. Traditional detection techniques such as the ELISA are generally limited to a single antigen and are therefore constrained to addressing only a small set of research hypotheses. In addition, most antibody detection assays are based on serum which is only one of a range of tissues in which these molecules mediate their effector functions. Except for blood-borne pathogens, antibodies exert most of their functional roles in tissue, including the mucosal surfaces of the respiratory, gastrointestinal and urinary tracts^5,6^. Protein microarray-based immunoassays are emerging technologies that permit the multiplexed detection of antibodies against a variety of antigens^7^. These technologies combine the capacity for highly parallel miniaturisation that is inherent to microarrays to the highly specific interactions between antigens and antibodies to construct a detection system that is capable of quantifying antibodies against hundreds of antigens within a single assay^8^. Peptide microarray-based immunoassays on the other hand have been used less extensively in clinical research and unlike protein microarray-based immunoassays, they utilise single linear epitopes from immuno-dominant regions of a protein as antigens^9-11^. Apart from mimicking well-defined epitopes that are highly specific to a particular pathogen^9,12,13^, this approach provides additional advantages over protein-based arrays, including reduced antigen cost, relative technical simplicity of peptide synthesis and the enhanced shelf-life of peptide array chips. Using a library of previously validated linear epitopes, we developed a synthetic peptide-based microarray for the detection of antibodies against multiple infectious diseases in a single microarray chip. We optimised the workflow to allow for the simultaneous quantification of two antibody subclasses in both serum and secretions derived from the airway mucosa of infants and young children. Using the data generated from the chip, we examined the systemic and secretory antibody repertoire against forty one infectious diseases in the first year of life. We also used the data to determine the kinetics of maternal antibody decay during this period.

## Materials and Methods

### Study site and population

The study was conducted in Kilifi County, a rural community on the Kenyan coast. 38 children (age range: 25 days – 17.5months) attending two local healthcare facilities (Kilifi County Hospital and the Pingilikani dispensary) with different infections were recruited into the study. Serum and nasopharyngeal and oropharyngeal swabs were collected from each child using previously described methods14. Written informed consent was obtained from the parents and legal guardians of all participants prior to recruitment. Ethical clearance for the conduct of the study was provided by the Kenya Medical Research Institute’s Scientific, Ethical and Research Unit.

### Selection of linear epitopes for the development of a peptide microarray immunoassay

We used the <u>Immune Epitope Database^15^</u> (IEDB) to construct a library of forty one experimentally-validated linear epitopes from a selection of infectious disease pathogens with moderate to high prevalence within the local population. Epitope sequences were sourced from previously published literature, where they had been independently validated using specific disease cohorts of naturally or experimentally-infected subjects. Two distinct epitopes were selected for each pathogen, with the epitope pairs in many cases being derived from different proteins. Details of the epitopes that were selected including source protein, amino acid sequence and source publication are outlined in table S1. The 82 linear epitopes (two for each of the 41 pathogens) were expressed as synthetic peptides by a commercial service provider (GenScript) with a free amino acid conjugated to each peptide.

### Development of peptide microarray chip

Lyophilised peptides were dissolved in a printing buffer consisting of 70% dimethyl sulfoxide, 5% glycerol and 25% sodium acetate (pH adjusted to 4.5). 50μl of the peptide solution was loaded onto the wells of a 384-well plate prior to printing. Peptides that did not fully dissolve were sonicated for 5 minutes and a further attempt to re-dissolve them was made. Functionalized 2D-epoxy glass slides (PolyAn, Germany) onto which the peptides were to be printed, were first dried on a heating block at 100oC for 10 minutes to remove residual moisture. Peptides were printed onto the epoxy slides using a non-contact inkjet microarray printer (Arrayjet, Scotland). Two peptide droplets of 100 picolitres each were printed onto the dried functionalized epoxy glass slides at a temperature of 20°C and 50% humidity. In addition to the peptides, the following control antigens were directly printed onto the slides: (i) an unlabelled capture antibody specific to the human IgG heavy chain and (ii) antibodies conjugated to fluorescent reporter labels (Alexa-Fluor 647 and Alexa Fluor 555: these antibodies were directly printed onto the slides in a pre-specified pattern as orientation landmarks). The capture antibody bound to all IgG molecules in the sample regardless of antigen specificity and provided process quality information on the printing and signal acquisition processes. In addition, negative control spots consisting of printing buffer were printed on each mini-array in order to identify non-specific antibody binding within individual mini-arrays. Printed slides were initially left to dry for one hour within the printing chamber and then at 25oC for 2 days in a dehumidified chamber. After drying, slides were then loaded onto 24-chamber hybridization cassettes and firmly locked into position. The slides were initially washed two times with PBS containing Tween-20 and then twice with only PBS. Blocking was done for one hour using 5% BSA in SSC buffer (0.25% SDS and 4.5% NaCl, adjusted to pH 7.0).

In order to detect mucosal antibodies in airway samples, transport media (UTM, Copan diagnostics) into which nasopharyngeal and oropharyngeal swabs (Copan diagnostics) had been eluted, were spun at 14000 rpm for 5 minutes in order to pellet cells and particulate debris. Supernatants were then centrifuged at 14000 rpm for 10 minutes in 3K Amicon centrifugal devices (Merck). The resulting concentrate was supplemented with an equal volume of 5% BSA in PBS containing Tween-20. A 50μl volume of each processed sample was then added to a single chamber of the hybridisation cassette and incubated for 4 hours at 4°C while shaking. For detection of serum antibodies, serum samples were diluted at 1:100 ratio in PBS and a 50μl volume added to the hybridisation cassette and incubated in the same manner as airway mucosa samples. After incubation, slides were washed as before. 1:500 dilutions of secondary antibodies - goat anti human IgA conjugated to Alexa-fluor 555 (Southern biotech) and goat anti-human IgG conjugated to Alexa flour 647 (Invitrogen) - were mixed together and then added onto the slides, which were left to incubate for 1 hour at 25oC. After incubation, slides were washed as above, disassembled from the cassettes, rinsed using Milli-Q water (Millipore) and spun dry for 5 minutes using a slide spinner. Scanning of the peptide microarray slides was done using a GenePix 4300A microarray scanner at wavelength of 635 nm and 532 nm at PMT levels of 500 and 100% power at 10μm resolution. Images were saved electronically in TIFF format and exported in JPEG format for visualization. Image analysis utilized Genepix Array List (GAL) files to extract quantitative values from spots. The quantitative values from the microarray scanner were expressed as background-corrected median fluorescent intensities (MFI). Correction for non-specific background signals was done automatically by the microarray scanner software. Scan results were then exported as tab separated value (tsv) files for further analysis.

### Data analysis

Data were analysed in the R programming environment16. Since each peptide was printed in duplicate, mean MFI values were calculated for each duplicate pair prior to analysis. For each pathogen, two distinct peptides were synthesised and printed onto the chip. Peptide-specific MFIs were compared between peptide pairs from the same pathogen and the peptide that exhibited the higher MFI signal was selected for further analysis. Loess curve-fitting was used to visualise the age-related antibody kinetics of a selection of antigens on the peptide microarray chip.

### Results

A synthetic peptide-based microarray for assaying IgA and IgG immunoglobulin subclasses against a range of infections in serum and airway-derived mucosal samples was developed successfully. Figure 1a shows an example slide with 24 mini-arrays that was used to simultaneously assay for IgG and IgA immunoglobulin subclasses in a set of 24 patient and control samples. The performance of the peptide microarray chip was evaluated by examining the expression levels of control spots. After processing the slides with patient samples, we compared the distribution of median fluorescence intensities of negative control spots (where buffer was printed) and positive control spots (where a capture antibody specific to the heavy chain of human IgG was printed). Almost all negative control spots had MFIs of 0 (geometric mean MFI=1.3), with a few spots having MFIs >10. Samples in mini-arrays with high negative control values were removed from further analysis. In contrast, all positive control spots resulted in high MFI values (geometric mean MFI = 32,359), with the majority of spots having mean MFI >10,000 (figure 1b). The coefficient of variation for the positive control spots was 7.6%.

**Figure 1:**
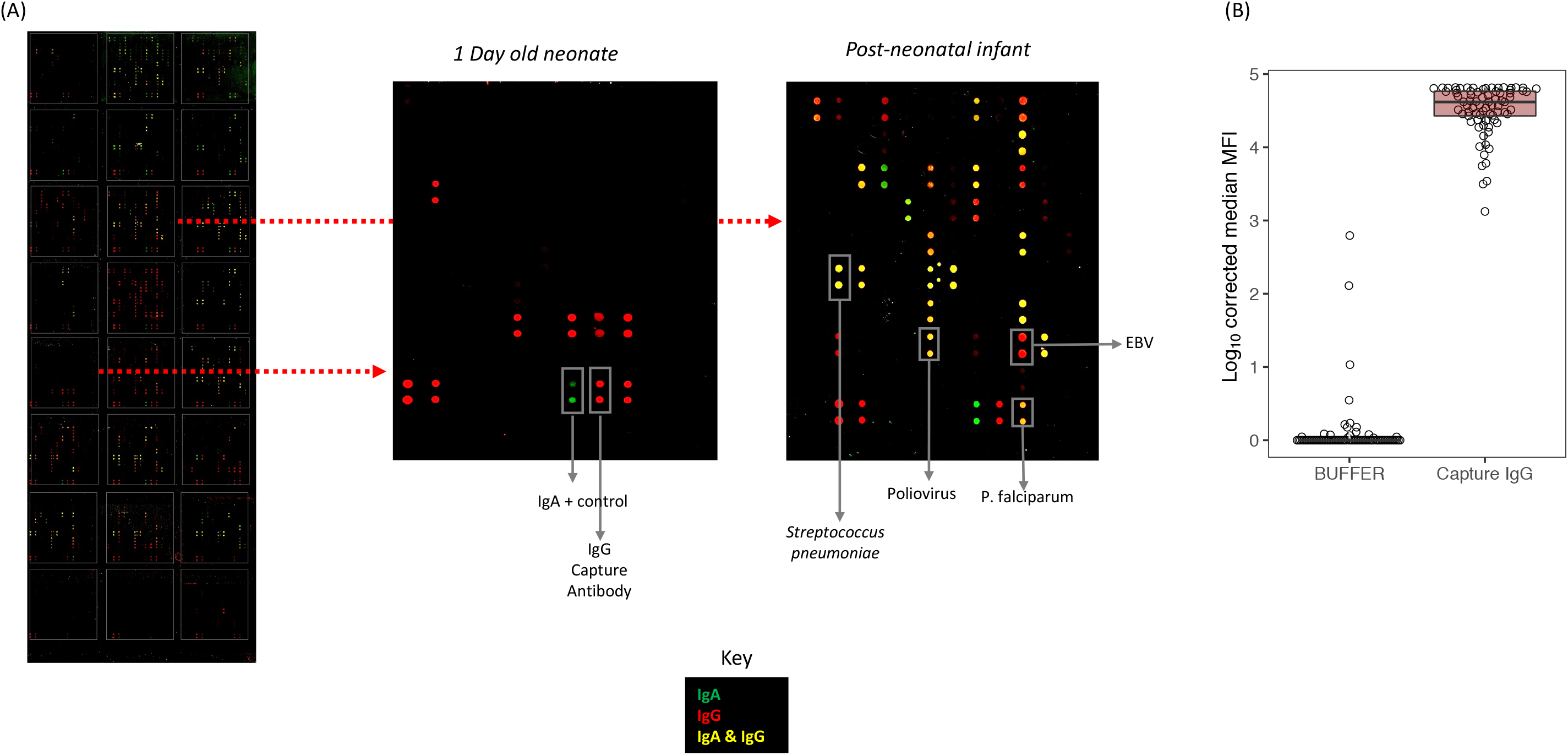
**(A)** An example slide in which immunoglobulins in sera from different children were assayed. Each microarray slide comprised 24 mini-arrays (shown using the white boxes). Two example mini-arrays are enlarged: in the first mini-array a sample obtained from a one-day old neonate was analysed while in the second, serum from a two-month old infant was analysed. On each mini-array, peptide epitopes were printed and serum/mucosal samples incubated. Antibodies in patient samples that bound to peptide epitopes were identified using a cocktail of two detection antibodies: anti human IgG conjugated to Alex Fluor (AF) 647 (red fluorescent signal) and anti-human IgA conjugated to AF-555(green fluorescence). Spots with a yellow hue indicate antigens which were bound by both IgA and IgG in patient samples. Each peptide epitope was printed in duplicate: the locations of a selection of duplicate antigen pairs are shown. **(B)** The distribution of fluorescence signals from control spots. Each mini-array contained a set of positive control spots (comprising of an anti-human IgG that bound to all IgG immunoglobulins in the patient sample irrespective of antigenic specificity) and negative control spots (microarray printing buffer). Almost all negative control spots did not yield a fluorescent signal - i.e. median fluorescent intensity (MFI)=0. The positive control spots on the other hand yielded much higher MFIs, with most spots resulting in MFIs >10,000.

Next we examined antigen-specific antibodies against all the antigens on the peptide array chip. This analysis was stratified by sample type (serum and airway mucosa) and immunoglobulin subclass (IgA and IgG). The most highly recognised antigens by serum IgG were Epstein Barr virus (EBVI), poliovirus (POLV), plasmodium falciparum (PFAL), *Streptococcus pneumoniae* (SPNE), Varicella Zoster (VZOS) and herpesvirus 1 & 2 (HV12), with geometric mean MFIs ranging from 15,189 (log_10_ 4.18) for EBVI to 42 (log_10_ 1.62 for HV12) – figure 2a. Apart from having the highest mean MFIs, these antigens were also the most frequently recognised, with the proportion of children who had detectable (MFI >0) serum IgG ranging from 100% (38/38) for EBVI to 58%(22/38) for HV12 (figure 2e). Mucosal IgG from the upper airway showed similar patterns of recognition; the six antigens with the highest mean IgG MFI levels in mucosal samples were EBVI (geometric mean MFI: 1,905), SPNE (geometric mean MFI: 953), PFAL(geometric mean MFI: 650), POLV(geometric mean MFI: 255) and VZOS (geometric mean MFI: 27) – figure 2b. Of these antigens, the most frequently recognised in mucosal secretions was SPNE, to which 95% (36/38) of the study population had detectable IgG, while VZOS was the least frequently detected was VZOS, with 60% (23/38) of the study population having detectable IgG – figure 2f. With respect to IgA, a similar profile of antigen recognition was observed in both serum and the airway mucosa. In serum, the three antigens with the highest levels of IgA reactivity were EBVI (geometric mean MFI: 230), SPNE(geometric mean MFI: 219) and VZOS (geometric mean MFI: 50) while in the airway mucosa, SPNE(geometric mean MFI: 93), PFAL(geometric mean MFI: 70) and POLV (geometric mean MFI: 13) had the highest mean IgA levels – (figure 2c & d).

**Figure 2:**
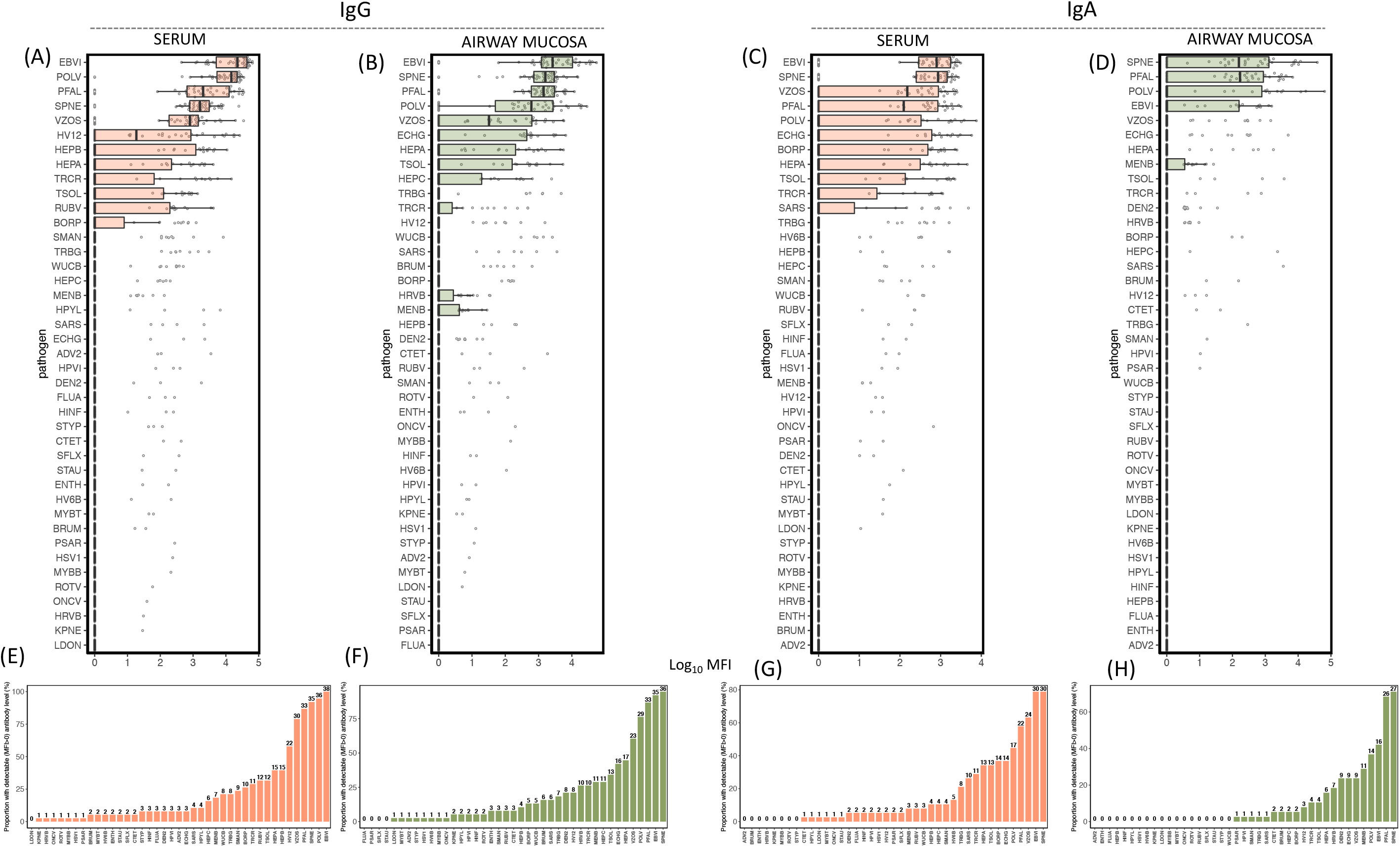
Immunoglobulin subclass-specific responses in serum and the airway mucosa **(A-D)** Distribution of IgG and IgA antigen specific MFIs in serum and mucosal samples. Antigens are ordered by immunoglobulin expression, with the antigens with highest mean MFIs occurring at the top of respective groups. The top five antigens by mean signal intensity in both samples were EBVI (Epstein-Barr virus), POLV (poliovirus), PFAL (plasmodium falciparum), SPNE (*Streptococcus pneumoniae*) and VZOS (varicella zoster). (E-F) The proportion of the study population with detectable (i.e. MFI>0) serum and mucosal antibodies against all antigens on the chip are shown. The numbers above each bar indicate the number of children who had detectable antibody levels for a particular antigen. Colour codes: orange – serum, green – airway mucosa.

To visualise global patterns of antigen recognition within the study population, we compiled antibody reactivity profiles at the individual level for all antigens on the peptide array chip (figure 3). For both IgG and IgA we observed broad heterogeneity in the antigens that were recognised by each individual in serum and in the airway mucosa. Each individual had different combinations of antigens that were recognised, although for certain antigens, there was near-universal recognition across the study population. With a few exceptions, most study participants had detectable IgG against EBVI, SPNE, PFAL and POLV in serum and airway mucosa samples (figure 3a-d). For some antigens, IgG recognition occurred more frequently in serum compared to the airway mucosa and vice versa. For example, echinococcus granulosus (ECHG)-specific mucosal IgG was detected in 42%(16/38) of study participants, while only 8%(3/38) had detectable IgG levels against this antigen in serum – (figures 2e & f and figures 3a & b). In contrast, rubella virus (RUBV) and HV12-specific IgG were detected more frequently in serum compared to the airway mucosa; 32%(12/38) and 39%(15/38) of the study population had detectable serum IgG to RUBV and HV12 respectively compared to 8%(3/38) and 21%(8/38) for whom RUBV andHV12 antibodies were detectable in airway mucosa samples (figures 2e & f and figures 3a & b). Some antigens were only minimally recognised (i.e. recognised by<5% of study population) in both serum and the airway mucosa; for example, low levels of antibody reactivity were observed against influenza A (FLUA), Leishmania donovani (LDON) and onchocerca volvulus (ONCV) where <5% of the study population had antigen-specific IgG or IgA (figures 2 & 3). The combinations of antigens that were recognised by IgA were generally similar to those recognised by IgG in both serum and the airway mucosa. In cases where antigens like SPNE and PFAL exhibited a high level of systemic and mucosal IgG reactivity, corresponding high IgA levels were observed in both serum and airway mucosa samples (figure 3).

**Figure 3:**
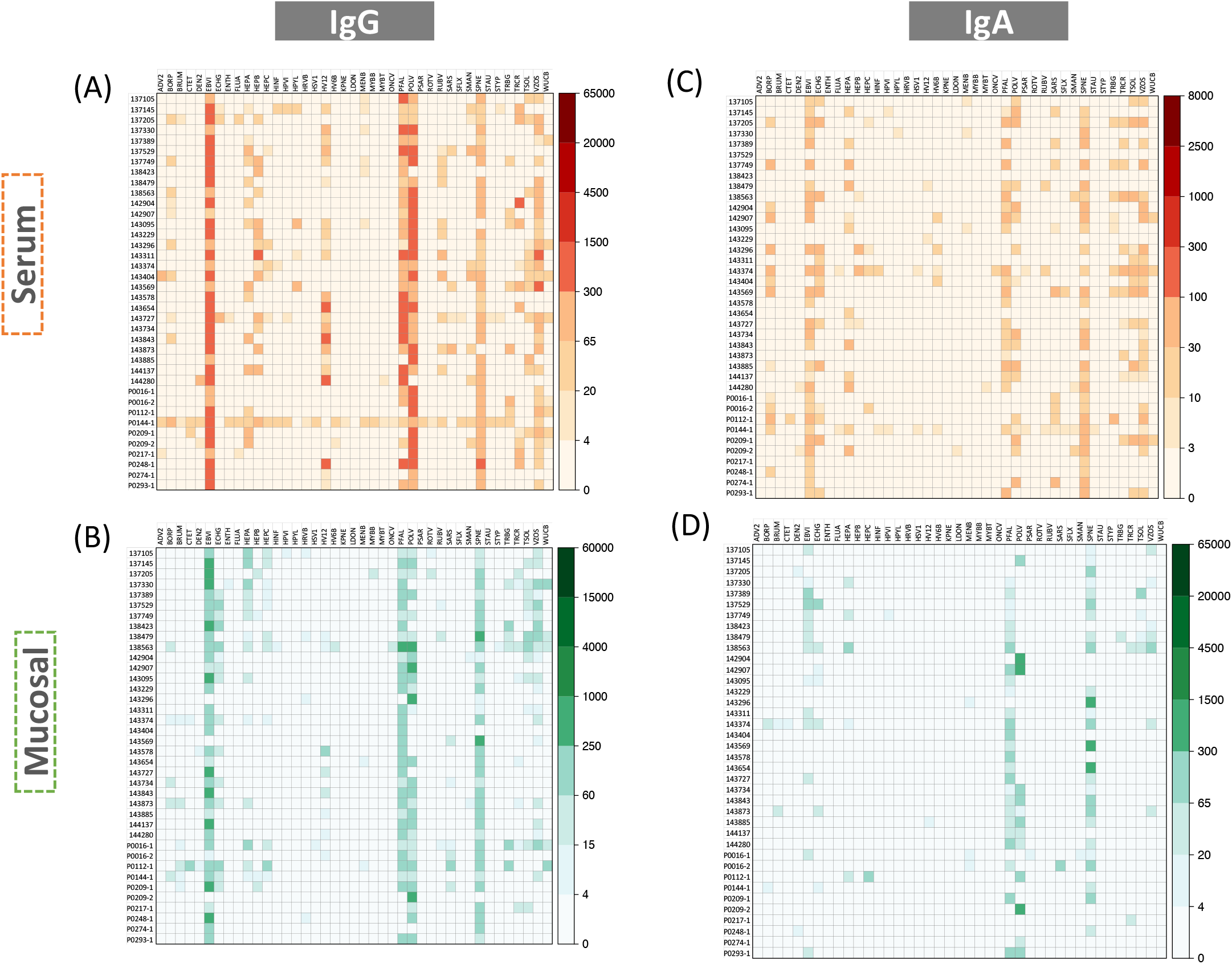
The combination of antigens recognised by different immunoglobulin classes in serum and airway mucosa are shown for each study participant. **(A-B)** Antigen specific IgG in serum and the airway mucosa are compared. Colour intensities correspond to the MFI signal intensity for a particular antigen. Within the study population, there was a broad overlap in the antigens that were detected in serum and the airway mucosa. **(C-D)** shows a similar comparison with respect to IgA.

Finally, we used the data obtained from the peptide microarray chip to examine the decay of antigen-specific maternal antibody in the study population. For this analysis, we focussed on the five antigens whose antibodies were most frequently detected in both serum and the airway mucosa: EBVI, POLV, SPNE, PFAL and VZOS. With the exception of SPNE, there was a general decline in serum IgG in the first six months of life (figure 4a). In the case of SPNE, IgG appeared to trend upwards in the first few months of life, peaking at around month six and maintaining this peak level thereafter. IgG levels in the airway mucosa showed a similar trend for all antigens except POLV: mucosa IgG specific to EBVI, SPNE, PFAL and VZOS declined sharply in the first six months of life and with the exception of SPNE, maintained this downward trajectory throughout the first 12 months of life (figure 4b). The dynamics of mucosal POLV IgG exhibited a markedly different trend: unlike the other antigens, there was a general rise in the level of POLV-specific mucosal IgG in the first six months of life (figure 4b, panel 2) followed by a later decline.

**Figure 4:**
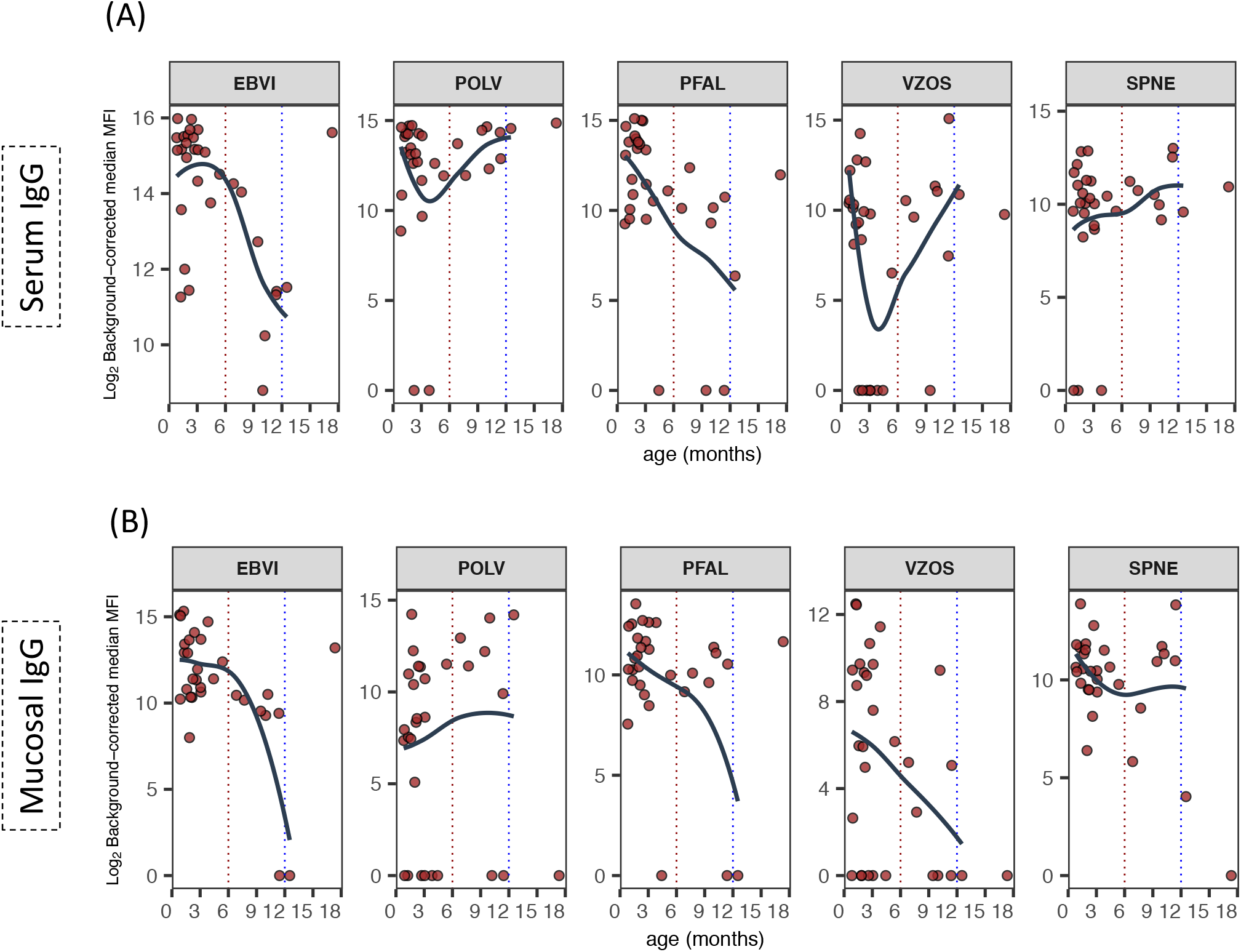
The kinetics of maternal antibody decay were evaluated for five antigens on the peptide microarray chip. Changes in the levels of antigen-specific IgG in **(A)** serum and **(B)** airway mucosa, were analysed using loess curve fitting. For all antigens except SPNE, there was evidence of serum IgG decline in the first three months of life. In the airway mucosa similar changes in antigen specific IgG were observed. The dashed red lines indicate the 6-month age time point while the blue dashed line indicates the 12-month time point.

### Discussion

We report the development of a synthetic peptide-based microarray for the measurement of antibody levels against forty-one infectious disease pathogens in serum and airway mucosal samples. The data from the microarray were used to explore the systemic and mucosal antibody repertoire against different infectious disease antigens in the first 18 months of life. Our data showed that during this period, most children had high levels of antibody against antigens derived from five infections: Epstein-Barr virus (EBV), poliovirus, *Streptococcus pneumoniae*, plasmodium falciparum and varicella zoster in both serum and the airway mucosa. In the first few months of life, it can be assumed that most of these antibodies are maternally-derived. The maternal antibody repertoire – delivered to the infant via breast milk or by trans-placental transfer – reflects lifelong maternal exposure to infectious disease and presents a snapshot of the locally prevalent infectious diseases^17^. In this study, some of the infections that were most frequently recognised within the study population have been previously documented to cause a very high disease burden within the east African coastal region in which the study recruits reside. For example, plasmodium falciparum, which causes malaria, has traditionally been endemic on the Kenyan coast and has been associated with a considerable annual morbidity and mortality burden over many decades^18-20^. In areas of comparable endemicity, approximately 40% of children under the age of five years have been reported to have circulating antibody against the parasite^21^, suggesting that women of child bearing age have high levels of circulating IgG available for transplacental transfer to foetal circulation. In the case of EBV, there is strong evidence from different parts of the world that infection is generally acquired within the first few years of life and that up to one in three children are EBV-seropositive by the age of five years^22-25^. This early exposure profile suggests that the widespread recognition of EBV antigens in this study could be traced to a combination factors including responses to natural infection as well as passively acquired maternal antibody as a result of long-term maternal exposure. In addition to EBV, we also observed an abundance of serum and mucosal antibody against the *Streptococcus pneumoniae*, a common mucosal pathobiont that normally colonises the upper respiratory tract in early life^26^ and is also the cause of pneumococcal pneumonia. Previous studies in Kilifi have shown that pneumonia is the biggest cause of childhood admissions^27^ and that pneumococcal pneumonia is responsible for about half of those admissions^28^. In addition to these pathogens, we also observed high antibody levels against poliovirus across the study population. We speculate that in addition to transplacental transfer, the poliovirus specific antibodies that were identified in the study population were as a result of vaccination. In Kenya, the oral polio vaccine (OPV) is administered to all infants at birth, 10 weeks and 14 weeks of age. This means that at the time of presentation to hospital, children in this study would have received at least one OPV dose and for a few of the children, all three doses. Taken together, the data presented are largely consistent with ancillary data on natural exposure, vaccination and maternal antibody transfer and suggests that the platform can be reliably used for more extensive serological analysis in future studies. In this study, we only included a limited number of antigens to provide initial proof-of-concept data. Due to its capacity for additional scalability, this platform can be expanded to incorporate and even broader array of infectious disease targets - including emerging threats such as ZIKA, Ebola and MERS - allowing a for more comprehensive profiling of humoral immunity in children.

We found evidence that antibodies against some antigens were differentially distributed between the serum and the airway mucosa. For example antibodies against echinococcus granulosus were detected more frequently in mucosal samples compared to serum. This difference might be attributed to the fact that echinococcus granulosus is a mucosal pathogen, and the host immune response against it is likely to be co-ordinated by immune cells resident in mucosal tissue. Animal studies have shown that experimental infection results in the production of high levels of anti echinococcus granulosus antibodies that can be detected in mucosal samples - including saliva - but not in serum^29^. Contrary to our expectations, some antigens that were derived from highly prevalent infections were detected at very low frequencies within the study population. For example, antibodies against influenza A (FLUA), a highly transmissible respiratory virus, were only detected sporadically. One possible explanation for this unexpected observation is antigenic mismatching between the assay antigen and that of locally circulating influenza A strains. The influenza A epitopes in this work were derived from the hemagglutinin (HA) protein, a highly variable protein that is characterised by temporal antigenic drifts in response to a build-up of population immunity^30^. Although we did not genetically characterise locally circulating influenza A strains over the course of the study, it unlikely that the HA sequences of these viruses would be identical to the HA antigen used on the microarray chip^31^ that circulated in the United States almost ten years earlier. Using data from the microarray, we also examined the age-related dynamics of antibodies specific to the five most frequently detected infections: EBV, poliovirus, *S. pneumoniae*, v. zoster and P. falciparum. With the exception of the pneumococcus, IgG levels in serum and the airway mucosa declined rapidly in the first three months of life, indicating the passive decay of maternally derived antibodies as has been extensively documented for numerous other infections^3,32,33^. In the case of the pneumococcus, we observed a general increase in the level of serum IgG throughout the first year of life. Previous studies have demonstrated that colonisation of the upper airway by the pneumococcus, tends to occur at a very early age. Studies in African infants have shown that over a third of all neonates are colonized with the pneumococcus within the first 28 days of life^34^. This early exposure profile might explain our failure to observe a decline in maternal pneumococcal IgG, since this decline would have been offset by the concurrent acquisition of natural antibody against colonising pneumococci. In sum, the observed kinetics of maternal antibodies in the first few months of life are consistent with expectations and largely align with previous observations. Apart from profiling antibody immunity in early life, the peptide microarray chip we’ve developed can be also be leveraged for different clinical and research applications including seroepidemiological studies. Previous studies have assayed antibodies to numerous infections as a proxy marker of disease exposure, and the resulting serology data have used to estimate disease burden^35-39^. The productive output of such seroepidemiological surveys can vastly be increased if in addition to the target antigen other related antigens are concurrently assayed, thereby providing a broader picture of disease interactions.

